# Multi-omics Profiling Reveals an NF-κB-Driven Anti-apoptotic Network Underlying Resistance to Oncolytic VSV in Prostate Cancer Cells

**DOI:** 10.64898/2026.06.24.734137

**Authors:** Alaa A Abdelmageed, Stephen Dewhurst, Maureen C Ferran

**Affiliations:** Biomedical Genetics and Genomics Program, University of Rochester School of Medicine and Dentistry, Rochester, NY 14642, USA; Department of Biomedical Genetics, University of Rochester School of Medicine and Dentistry, Rochester, NY 14642, USA; Gosnell School for Life Sciences, Rochester Institute of Technology, Rochester, NY 14623, USA

## Abstract

The therapeutic efficacy of oncolytic viruses is often limited by the presence of tumor cells that resist virus-mediated killing. Here, we investigated the molecular mechanisms underlying resistance to Vesicular Stomatitis Virus (VSV) in PC3 cells, an aggressive metastatic prostate cancer (PrCa) cell line, using the VSV-sensitive LNCaP cell line as a comparator. RNA sequencing revealed that, relative to untreated cells, VSV-infected PC3 cells upregulated both pro-apoptotic genes, including BIM, PUMA, and NOXA, and anti-apoptotic and antiviral genes, including A20 and RIG-I. In addition, genes associated with antiviral and pro-survival pathways, including NFκB and PI3K-Akt signaling, were more highly expressed in PC3 cells than in LNCaP cells. At baseline, PC3 cells also exhibited elevated expression of multiple pro-survival genes, including BCL-xL, MCL1, and CK2, compared with LNCaP cells. Complementary proteomic analyses identified enhanced activation of NFκB, PI3K-Akt, and MSK1 signaling in VSV-infected PC3 cells relative to infected LNCaP cells. Furthermore, pharmacological inhibition of BCL-2 family proteins or NFκB signaling restored sensitivity to VSV-induced cell death in PC3 cells. Collectively, these findings identify NFκB-centered pro-survival signaling networks as key contributors to the resistant phenotype of PC3 cells and suggest that combining oncolytic virotherapy with targeted inhibitors may improve therapeutic efficacy in resistant prostate cancers.

## Introduction

Oncolytic viruses are a novel yet powerful tool for cancer therapeutics. In recent years, VSV, a negative-sense RNA virus belonging to the family *Rhabdoviridae*, has gained attention as an exceptionally promising oncolytic virus [1,2]. VSV displays broad cell tropism, infecting nearly all mammalian cell types, yet lacks pre-existing immunity in humans [3]. Its inherit cancer tropism and cancer cell-specific lysis could be attributed to the defectiveness of the interferon (IFN) signaling many cancer cells harbor [4–6]. Despite its initial success in pre-clinical and some clinical studies, some tumors actively mount antiviral defenses, therefore becoming VSV-resistant [7,8]. While previous research claims the IFN pathway drives this resistance, the exact mechanisms remain poorly understood, especially in the context of prostate cancer [9,10]. As of 2026, it is estimated that prostate cancer is the most common cancer in men (<u>11</u>). If diagnosed early, its 5-year survival rate is nearly a 100%; however, this prognosis for metastatic and distant stages only applies to roughly 30% of cases, with castration-resistant types receiving the worst prognosis [12]. In this study, we use the two PrCa cell lines to model tumors that are either VSV-resistant or VSV-sensitive, respectively [13]. LNCaP cells were initially isolated from a supraclavicular lymph node adenocarcinoma metastasis. They express androgen receptor (AR) and therefore represent a less-aggressive PrCa that is responsive to androgen-deprivation (castration) therapy. Conversely, PC3 cells that were first isolated from a bone metastasis, resemble a more aggressive, castration-resistant PrCa and do not express AR [14]. When infecting the two cell types with VSV, an MOI of 10 is used for LNCaP, while an MOI of 50 is used for PC3 to establish a synchronous infection [15]. We have previously demonstrated that the NF-κB pathway is central to establishing the resistance to VSV-mediated cell death, with multiple steps being activated in the pathway in PC3 but not LNCaP cells [16]. Here, we took a comprehensive approach to investigating the underlying resistance mechanisms in PC3 cells. This was done through a multi-omic and experimental approach. RNA sequencing was used to understand the transcriptional layout in the two cell types, and their differential response to VSV infection. This was followed by a phospho-proteomic array of proteins involved in the NF-κB pathway, both directly and indirectly. Lastly, we performed a pharmacological blockage of anti-apoptotic genes downstream from the NF-κB pathway in PC3 cells, using both WT VSV and the M51R (R1) mutant commonly utilized in preclinical studies [17].

This research provides insights into the underlying mechanisms of resistance to VSV by aggressive PrCa, and an understanding of other pathways interacting with NF-κB to produce this phenotype. Several strategies to overcome resistance to VSV have been suggested in the past [18]. Such strategies, along with our proposed focus on anti-apoptotic factors, can improve the potential of VSV oncolytic therapy for PrCa.

## Methods

### Cells, Viruses, and Infections

The LNCaP and PC3 human PrCa cell lines were obtained from the American Type Culture Collection (ATCC) (CRL-1740 and CRL-1435, respectively) and were grown in RPMI-1640 containing 10% FBS (ATCC). The heat-resistant (HR-C) strain of the Indiana serotype of VSV was used as the WT virus. Mutant T1026R1 (R1), a temperature-stable revertant of T1026, was derived from the Indiana HR-C strain of VSV by chemical mutagenesis (supplied by C. P. Stanners). The R1 mutant, which harbors a single amino acid substitution at position 51 of the M protein (M51R), was used as a representative example of a virus strain that is attenuated relative to the WT virus—and thus has a more favorable safety profile for potential use as an oncolytic agent in humans [19]. All viruses were grown on Vero cells as described [20,21] in EMEM growth medium (ATCC). Cells were infected with each virus at multiplicities of infection (MOI) of 10 PFUs/cell for LNCaP and 50 PFUs/cell for PC3 cells. Viruses were allowed to adsorb to target cells in serum-free culture medium for 1 h at 37 °C, after which complete (serum-containing) culture medium was added to the cells.

### PrestoBlue Viability Assay

PC3 and LNCaP cells were seeded in Santa Cruz Biotech black frame/clear bottom cell culture 96-well plates (sc-204468). After 24 h, cells were infected with either WT VSV or the R1 mutant at an MOI of 0.1 or 1. At 24, 48 and 72 h post infection (hpi), supernatants were removed and 100 µL of master mix was added, composed of 10% PrestoBlue HS Cell Viability Reagent (Thermo Fisher Sci, Waltham MA, USA) in serum-free, phenol red-free RPMI medium. This viability reagent is reduced by viable cells and converted into a red fluorescent dye. Plates were then incubated in the dark at 37 °C for 40 min and a fluorescence reading was taken on a SpectraMax iD3 Plate Reader (Molecular Devices, San Jose CA, USA) using an Em/Ex 590/550 nm wavelength range.

### RNA Sequencing

Cells were grown in 30 mm tissue culture dishes coated with poly-D-lysine (PDL). Infection was carried out using VSV at an MOI of 50 and 10 PFU/cell for PC3 and LNCaP, respectively. At 6 hpi for LNCaP and 8 hpi for PC3 cells, cells were lysed in plates using RIPA lysis buffer (Thermo Scientific, Waltham MA, USA; RIPA Lysis and Extraction Buffer). Samples were sent to the University of Rochester Genomic Center for library preparation and sequencing. Analysis was performed on count data using the Bioconductor DESeq2 package (Version 1.44.0) to calculate differential expression (DE) levels. Contrasts were made on the DE data by comparing virus treatments to mock treatment for each cell line. Data normalization was performed using vst on the dds object and within the DESeq pipeline itself.

### BCL-2 family inhibitor drugs

Cells were treated with 5 µM of ABT-199, 5 µM of ABT-737, or 10 µM of IKK16 purchased from Med Chem Express, Monmouth Junction NJ, USA; cat#: HY-15531, HY-50907, HY-13687 (respectively) for 24, 48 and 72 h. 6 h prior to each timepoint, LNCaP cells were infected with an MOI of 10, while at 8 h prior to each timepoint, PC3 cells were infected with an MOI of 50. Following infection, samples were treated with PrestoBlue, and readout was measured as fluorescence with a SpectraMax iD3 Plate Reader (Molecular Devices).

### Proteomics Microarray

Cells were grown in 30 mm tissue culture dishes coated with poly-D-lysine (PDL). Infection was carried out using VSV at an MOI of 50 and 10 PFU/cell for PC3 and LNCaP, respectively. At 6 hpi for LNCaP and 8 hpi for PC3 cells, cells were lysed in plates using RIPA lysis buffer (Thermo Scientific, Waltham MA, USA; RIPA Lysis and Extraction Buffer). Samples were sent to the Full Moon Biosystems, Sunnyvale CA, USA for performing an array of 217 antibodies to detect expression of proteins interacting with the NF-κB pathway. Analysis was performed on array data using the Bioconductor limma package (Version 1.44.0) to calculate differential expression (DE) levels. Contrasts were made on the DE data by comparing virus treatments to mock treatment for each cell line.

### Statistical Analysis

Statistical analysis throughout this paper was performed in RStudio (Version R version 4.4.0 (2024-04-24) – “puppy cup”) using either the Student’s t test or the one-way ANOVA test and an asterisk indicates a significant difference (p < 0.05). Unless otherwise noted, results are expressed as means and error bars indicate the ±standard error of the means (SEM).

## Results

### PC3 cells respond to VSV infection and transcriptionally activate pro-apoptotic genes

To understand the mechanisms underlying VSV-resistance in PC3 cells, we performed a transcriptomic analysis of infected PC3 and LNCaP cells. Cells were infected with an MOI of 50 for PC3 cells and 10 for LNCaP to establish a synchronous infection (note that a higher MOI was necessary for PC3 cells, due to their greater resistance to VSV infection, as described in detail elsewhere) [15]. Principal component analysis (PCA) revealed that VSV infection induced substantial global transcriptional changes in both cell lines, with comparable separation between infected and uninfected samples. (Figure. 1.A).

**Figure 1.**
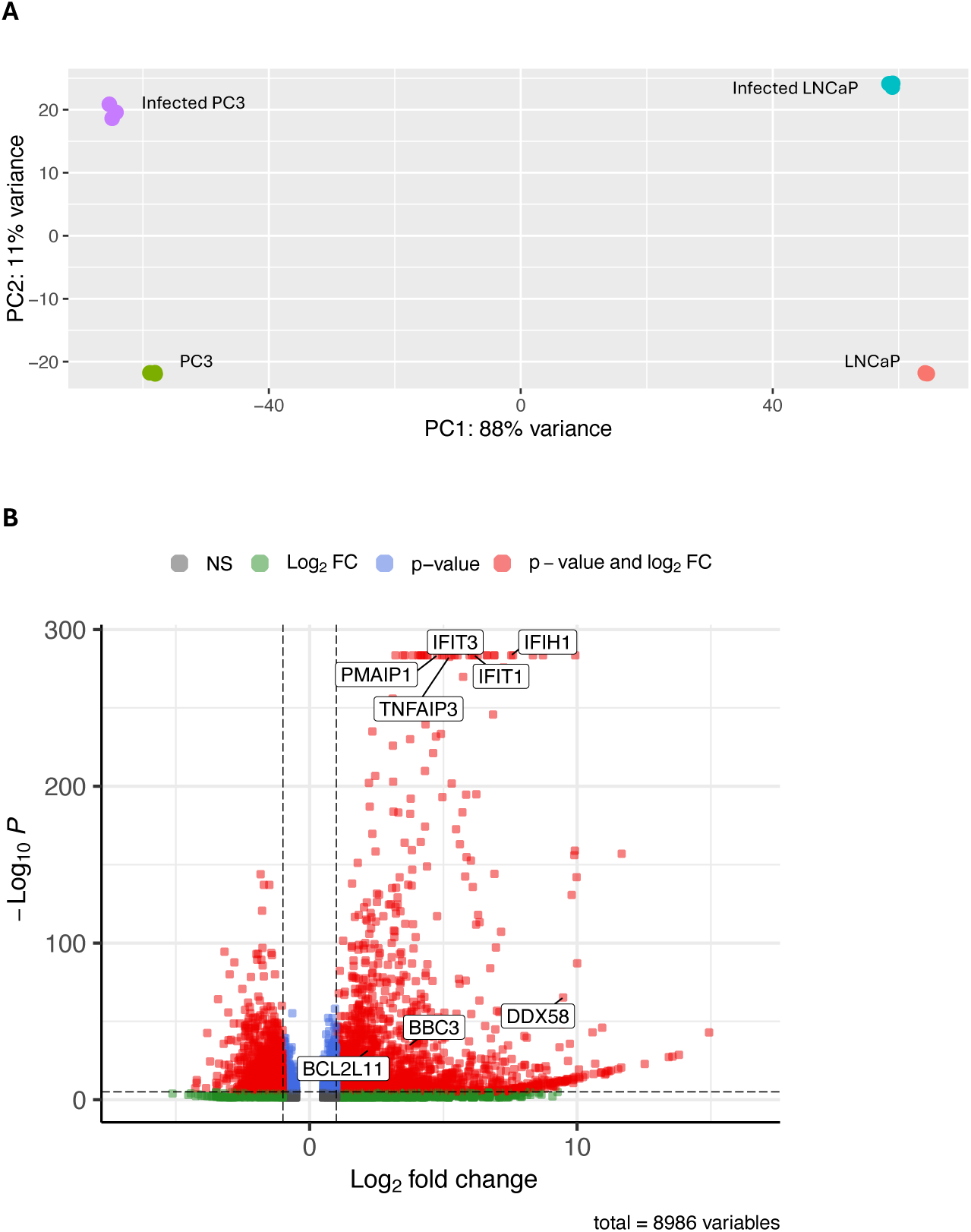
VSV infection results in transcriptional reprogramming in PC3 and LNCaP cells. A) PCA plot of all cellular mRNAs in infected PC3 and LNCaP cells, as compared to uninfected control cells. B) Scatter plot of differentially expressed genes in infected PC3 cells in comparison to the untreated cells. A cutoff of <0.05 for p-adjusted values was applied, and >|0.5| for log2FoldChange. Plot was created using the EnhancedVolcano (v.1.24.0) package in R.

VSV infection of the resistant PC3 cells induced the expression of several pro-apoptotic genes, including BCL2L1 (BIM), BBC3 (PUMA), and PMAIP1 (NOXA) (Figure. 1.B). Nevertheless, previous research has shown that PC3 cells resist killing by VSV as compared to LNCaP cells [16]. Consistent with this, VSV infection also upregulates many anti-apoptotic factors like TNFAIP3 (A20) – as well as antiviral genes such as DDX58 (RIG-I) and IFIT1&3 (viral RNA sensing) in PC3 cells. This simultaneous expression of both pro- and anti-apoptotic genes suggests that the outcome of viral infection (cell death vs. survival) in PC3 cells and in LNCaP cells is a reflection of the balance between these pro- and anti-apoptotic factors.

### PC3 cells express higher basal levels of anti-apoptotic/pro-survival genes

To understand the factors that contribute to the greater resistance of PC3 cells to VSV infection, as compared to LNCaP cells, we compared the baseline patterns of gene expression in both cell lines, in the absence of viral infection. PC3 cells expressed higher levels of many anti-apoptotic genes like BCL-xL of the BCL-2 family, BIRC2 (cIAP1), CK2, and MCL1 (Figure. 2.A-C, G). They also showed higher expression of NF-κB pathway components like p100 and RELB (Figure. 2.D, E). Notably, LNCaP cells expressed somewhat higher levels of the anti-apoptotic BCL-2 gene (Figure. 2.F). Overall, the more abundant baseline expression of anti-apoptotic/pro-survival genes in PC3 cells may explain, in part at least, the greater resistance of this cell line to VSV-mediated cell killing.

**Figure 2.**
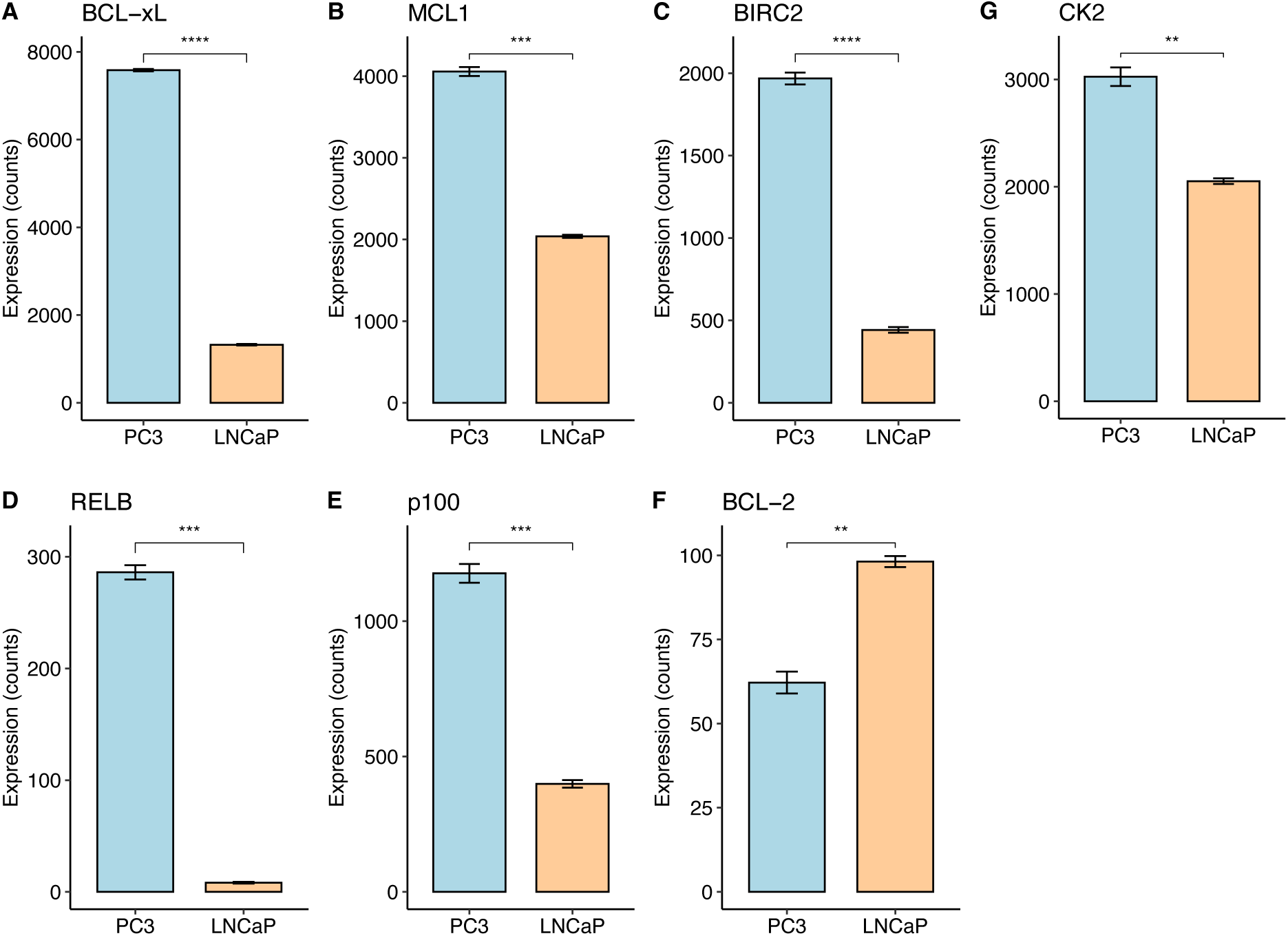
PC3 cells express higher baseline levels of many anti-apoptotic/pro-survival factors than LNCaP cells. RNA sequencing data showed higher expression of A) BCL-xL B) MCL1 C) BIRC2 D) RELB E) p100 and G) CK2 in PC3 cells at baseline, as compared to LNCaP cells. (*n* = 3, Mean ± SEM, **** p < 0.0001, *** p < 0.001, ** p < 0.01, * p < 0.05, Student’s *t*-test).

### Many pro-survival and antiviral pathways are enriched in infected PC3 cells

To investigate how VSV infection affects the gene expression portfolio of the two cell lines, overrepresentation analysis (ORA) was performed separately on genes differentially expressed following infection in PC3 and LNCaP cells relative to their respective uninfected controls. The ORA showed selective enrichment of p53, IL-17, Apoptosis, and MAPK signaling pathways in infected PC3 cells, as well as enrichment of the TNF, cytokine-cytokine receptor interaction, and NF-κB signaling pathways in both cell lines (Figure 3). We note that this analysis does not represent a direct comparison of infected PC3 and LNCaP cells, but rather, it is a side-by-side comparison of the response of each cell line to virus-infection (as compared to their uninfected baseline state).

**Figure 3:**
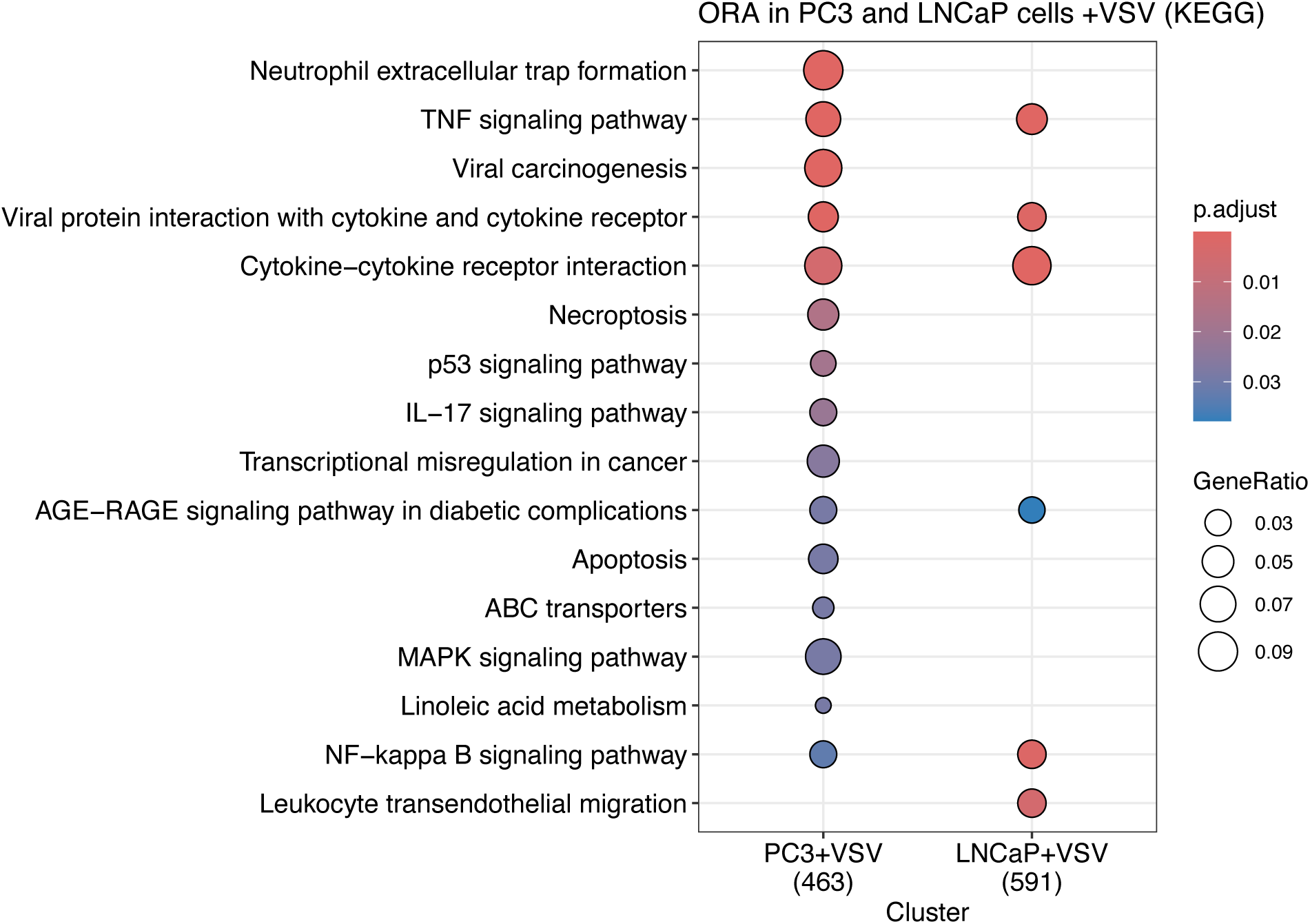
A subset of KEGG pathways, including IL-17 signaling, apoptosis and p53 signaling, are selectively enriched in infected PC3 cells versus LNCaP cells. Shown here is an overrepresentation analysis of enriched KEGG pathways in infected PC3 (left) and LNCaP (right) cells, as compared to their matched, uninfected counterparts. Gene lists were generated using a cutoff of <0.001 for p-adjusted values and >|2| for log2FoldChange. The BioConductor packages clusterProfiler (v. 4.20.0) and enricplot (v.1.32.0) were used to generate this plot in R.

### PC3 cells display stronger basal expression of antiviral and pro-survival pathways

To follow up on our ORA, we generated and examined more detailed expression maps of the enriched pathways to analyze gene expression signatures along the pathway. At baseline, PC3 cells showed higher expression of many pro-survival genes in the NF-κB pathway, including BCL2A1 and PLAU (Figure 4). Many genes in the PI3K-Akt pathway also displayed higher baseline expression in uninfected PC3 cells as compared to uninfected LNCaP cells (Figure S.1). Interestingly, PC3 cells expressed higher levels of many pro-inflammatory cytokines and chemokines in the TNF pathway, such as IL6 and CXCL1 (Figure S.2). In addition, many of the pro-survival genes in the apoptosis pathway were also more highly expressed in uninfected PC3 cells as compared to uninfected LNCaP cells (such as BIRC2, TRAF1, and BCL2A1; Figure S.3).

**Figure 4:**
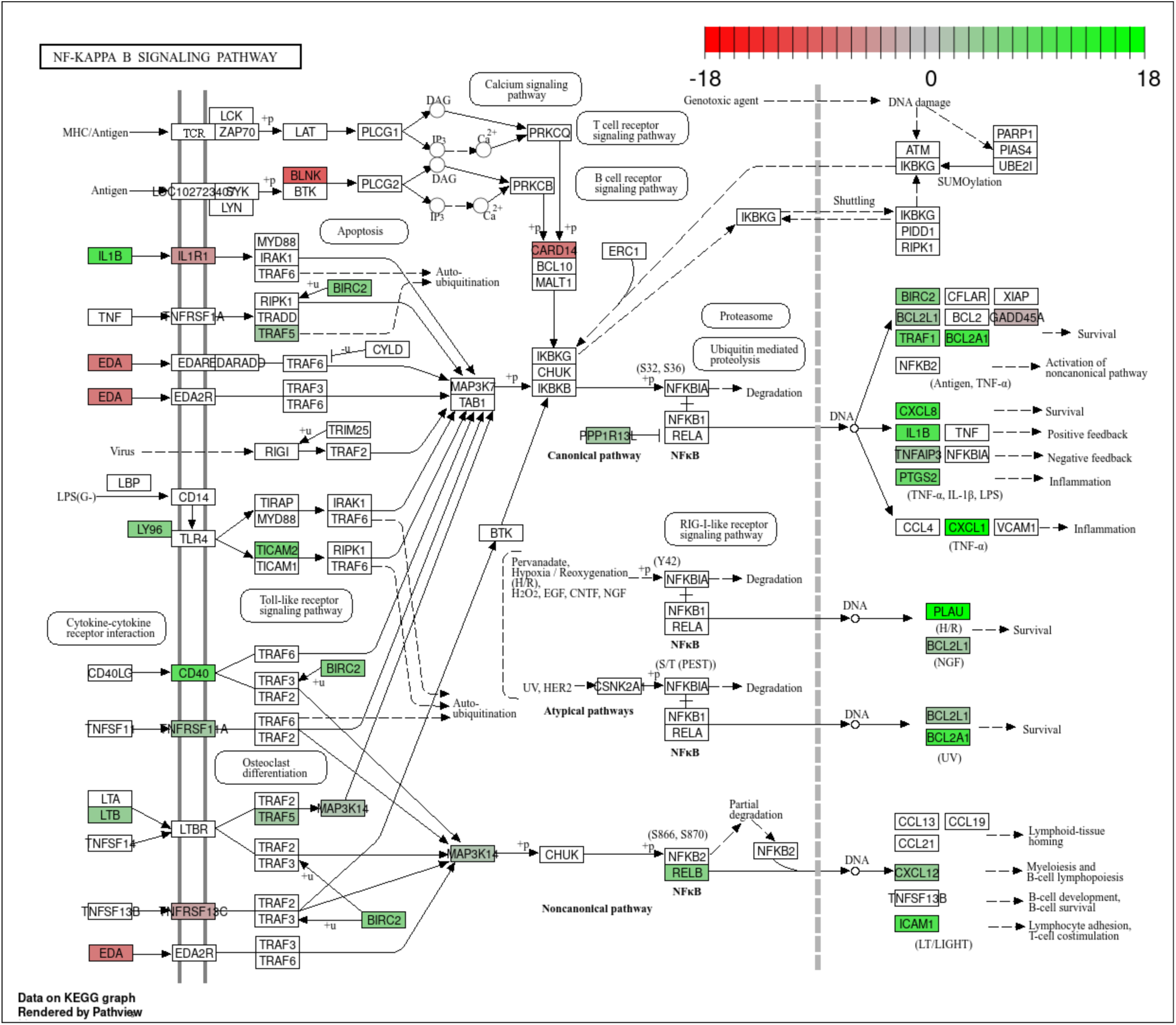
PC3 cells express higher levels of many pro-survival genes in the absence of viral infection, as compared to LNCaP cells. KEGG pathway enrichment analysis of the NF-κB pathway in uninfected PC3 cells as compared to LNCaP. The color scale reflects log2 Fold Changes (green = upregulated in PC3 cells, gray = unchanged, red = downregulated (vs. LNCaP cells)). The Bioconductor package pathview (v.1.52.0) was used to generate this figure in R.

### VSV infection sustains the higher expression of antiviral and pro-survival pathways in PC3 cells

Having dissected differences in key gene expression pathways in uninfected PC3 and LNCaP cells, we next turned our attention to their infected counterparts. To do this, we compared gene expression in VSV-infected PC3 and LNCaP cells across multiple KEGG pathways. PC3 cells sustained higher expression of the same genes presented in figure 4 (and Figures S1-S4) relative to LNCaP cells, including those in the NF-κB (Figure 5), PI3k-Akt, TNF, apoptosis, and prostate cancer pathways (Figures S5-S8). These findings indicate that many transcriptional differences observed at baseline were maintained following infection (and in some cases, even increased – see Figures S9, S10) following virus infection.

**Figure 5:**
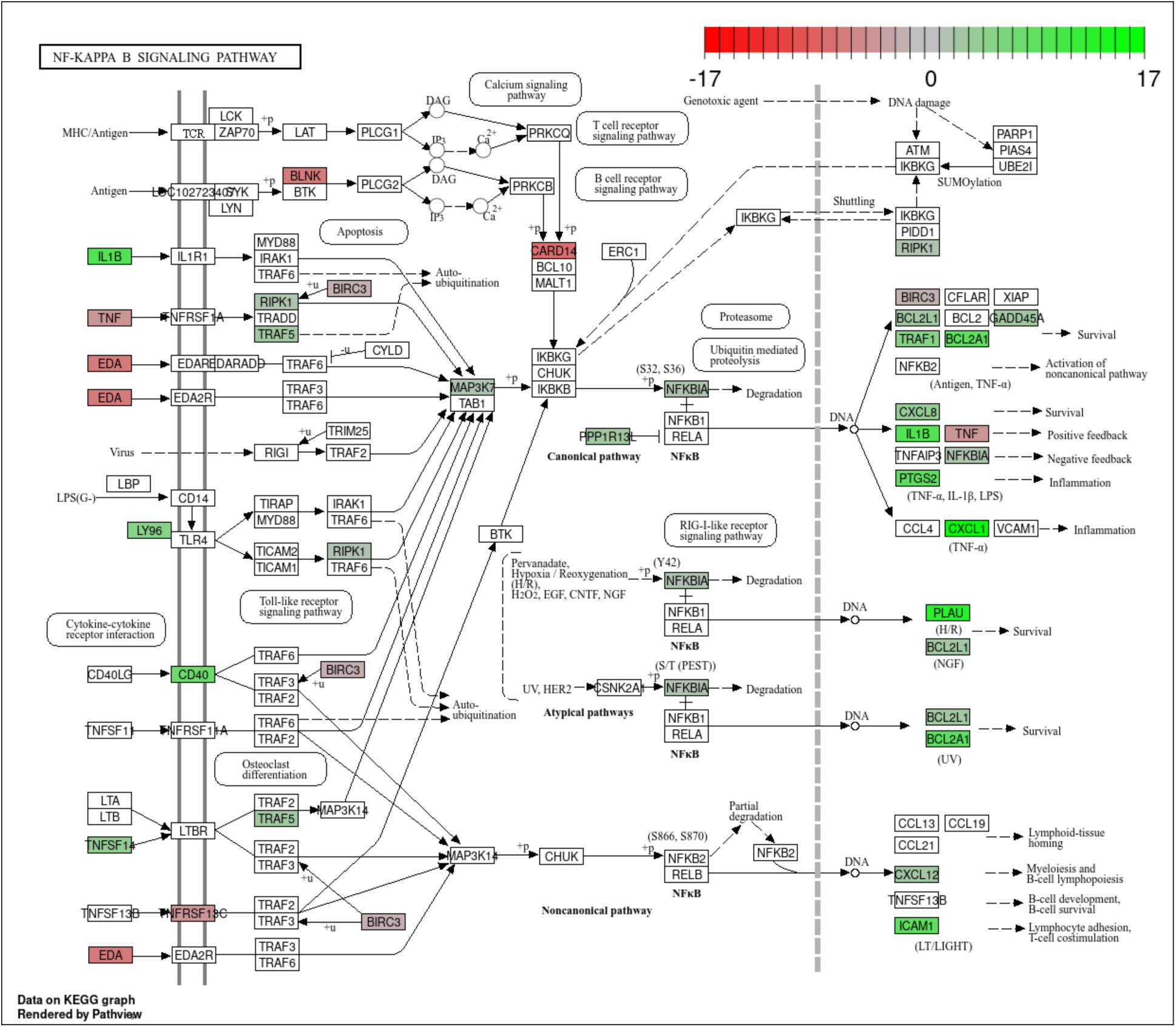
VSV infection causes upregulation of many pro-survival and antiviral genes in PC3 cells, as compared to LNCaP cells. KEGG NF-κB pathway enrichment analysis in VSV-infected PC3 cells compared to infected LNCaP cells. The color scale reflects log2 Fold Changes (green = upregulated in PC3 cells, gray = unchanged, red = downregulated).. The Bioconductor package pathview (v.1.52.0) was used to generate this figure in R.

### VSV infection results in activation of the NF-κB and Akt pathways in PC3 cells

One limitation of transcriptional analyses is that they cannot account for changes at the post-translational level. We therefore performed proteomics analysis on infected PC3 and LNCaP cells, by subjecting cell lysates to analysis using array of 217 total and phospho-specific antibodies directed against proteins in the NF-κB pathway (Fig. 3.6). As predicted by our transcriptional analyses, PKR was activated in infected PC3 cells (as reflected by an increase in phosphorylation at Thr 446) compared to their uninfected counterparts. This protein-based array analysis also revealed the increased upregulation (and activation) of proteins involved in both the canonical and non-canonical NF-κB pathways downstream from TNFR2 in PC3 cells, relative to LNCaP cells (as reflected by increased phosphorylation of key regulatory amino acids in multiple NF-κB family members; Fig. 3.6). Furthermore, proteins in the PI3K-Akt pathway were also activated in response to VSV infection in PC3 cells compared to LNCaP cells (as reflected also by increased phosphorylation of key regulatory residues; Fig. 3.6). Finally, IKK*α* and IKKβ were downregulated in PC3 cells following infection, but MSK1 was activated; the latter may reflect both a stress response to viral infection, and an additional contribution to NF-κB activation, possibly through phosphorylation by MSK1 (Figure 6).

**Figure 6:**
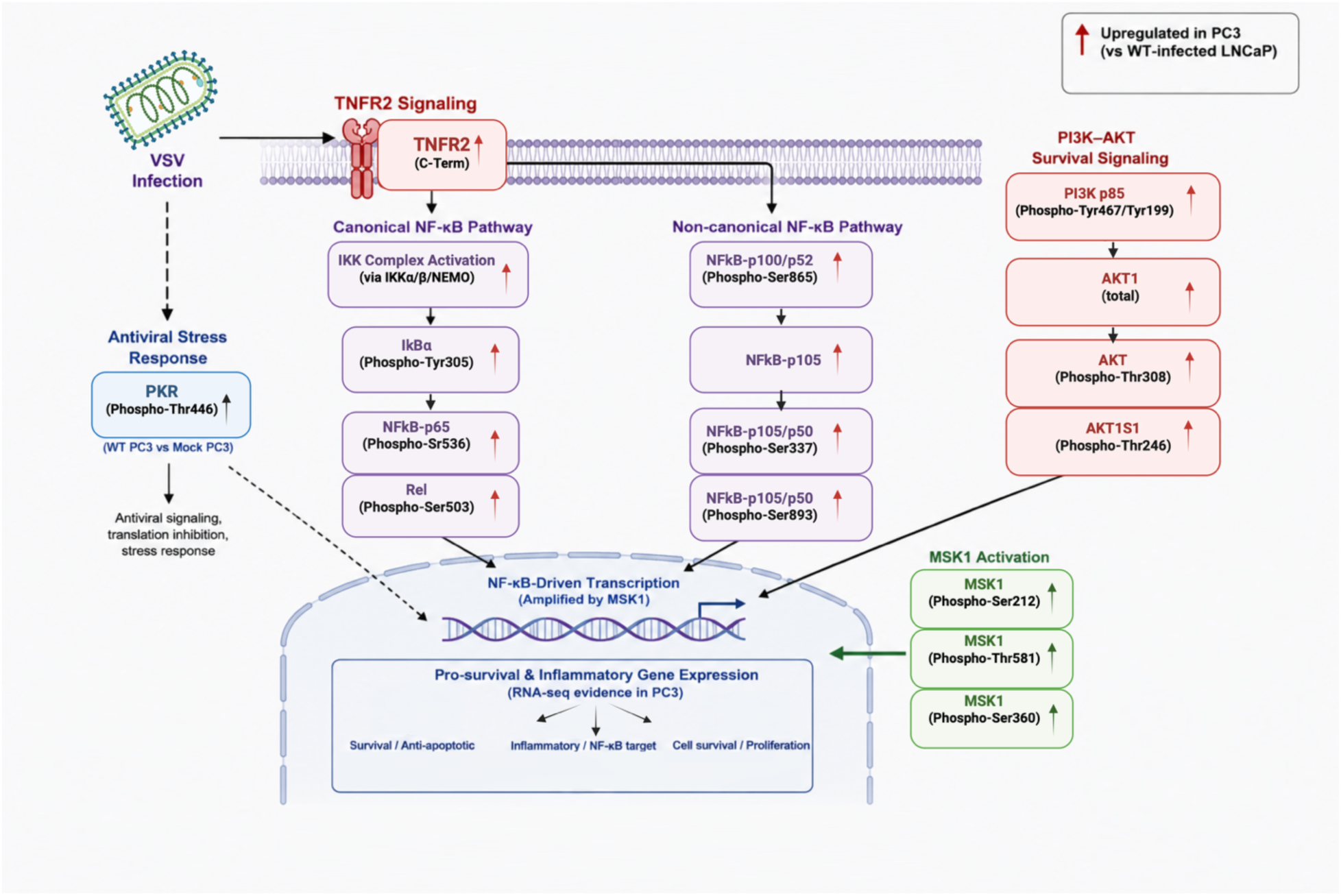
VSV infection of PC3 cells activates many proteins in the NF-κB and PI3K-Akt pathways. Cell lysates from infected PC3 and LNCaP cells were interrogated with a panel of 217 antibodies directed against proteins represented in the NF-κB and PI3K-Akt pathways (including phosphor-specific antibodies capable to detecting key post-translational modifications associated with pathway activation). To analyze these results, DE analysis was performed in R using the limma package (v.1.44.0). Filtration was performed using a cutoff of 0.05 for adjusted p-value and > |0.25| for log2 fold changes to account for slight changes in post-translational modifications.

### Pharmacological blocking of anti-apoptotic proteins reverses PC3 cells resistance to VSV

Our transcriptomic and proteomics data suggested possible underlying mechanisms of VSV-resistance in PC3 cells. To confirm the centrality of anti-apoptotic factors to cell fate, following virus infection, pharmacological inhibitors of BCL-2 family members were used to treat PC3 cells with and without VSV infection. For these experiments, we used ABT-199 (aka Venetoclax), which is an established chemotherapeutic drug that mainly targets BCL-2, and ABT-737, which targets multiple BCL-2 family members (including the anti-apoptotic genes BCL-2, BCL-xL, and BCL-w). As expected, neither drug treatment alone, nor viral infection alone, widespread cell death. However, viral infection of drug-treated PC3 cells caused overwhelming cell death (Figure 7). Similar findings were obtained in PC3 cells exposed to the NF-κB pathway inhibitor, IKK16 – resulting in cell death only in cells pretreated with IKK16 and then infected with VSV (Figure 7). Collectively, these findings support a role for anti-apoptotic and NF-κB signaling pathways in the resistance phenotype of PC3 cells

**Figure 7:**
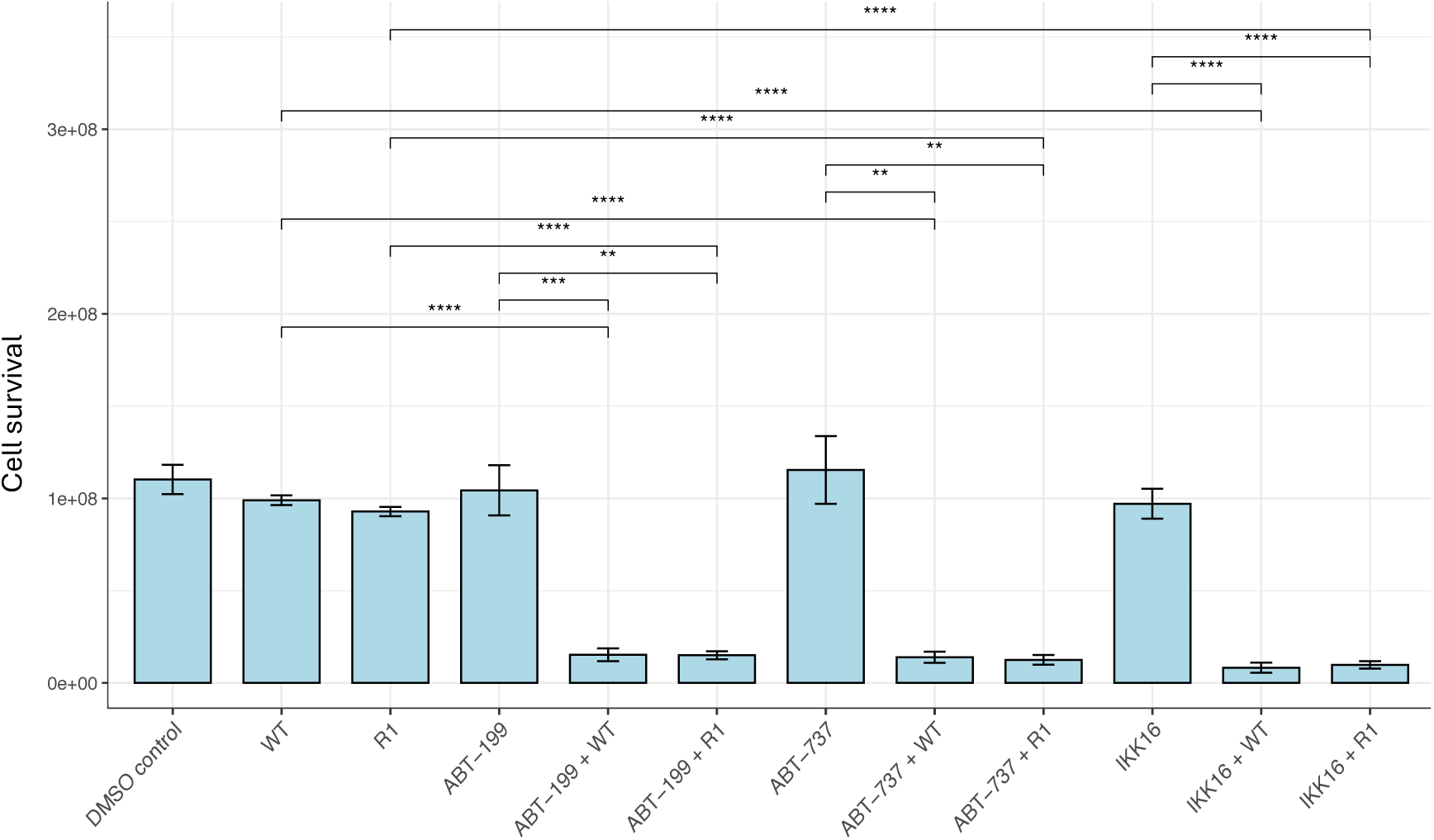
Inhibition of the anti-apoptotic BCL-2 and NF-κB pathways restores sensitivity to VSV infection in PC3 cellsPC3 cells were exposed to selective BCL-2 pathway inhibitors (ABT-199 and ABT-737) or a selective NF-κB pathway inhibitor (IKK16) either alone, or in the presence of VSV infection (at an MOI of 50, using either wild-type virus or the attenuated R1 mutant). Cell survival was then quantitated using the PrestoBlue cell viability assay at 24 post-treatment with each drug. Viral infection was initiated 8 h prior to analysis at each timepoint. Data represent the mean ± SEM of 3 biological replicates. **** *p* < 0.0001, *** *p* < 0.001, ** *p* < 0.01 (Student’s *t*-test).

Finally, we also tested whether the cells elicited a similar response when cells were exposed to the same drugs in the setting of infection with the R1 VSV mutant. This mutant virus harbors a mutation in M at position 51 and is a safer option to use in a clinical setting (as compared to WT VSV), with reduced potential for neurotoxicity. Co-treatments with the drugs and R1 resulted in a similar response to that of WT-drugs co-treatment, also producing a synergistic effect and leading to cell death in VSV-infected PC3 cells (Figure 7).

## Discussion

Conventional cancer therapies often suffer from limited tumor selectivity and can result in significant toxicity to healthy tissues. In contrast, oncolytic viruses offer a promising alternative because of their ability to selectively infect and kill malignant cells, while sparing healthy cells. However, subsets of tumor cells may resist virus-mediated killing and ultimately limit therapeutic efficacy. Therefore, understanding the molecular basis of this resistance is critical for the development of more effective oncolytic virus-based therapies.

In this study, we used the PC3 and LNCaP PrCa cell lines as representative models of VSV-resistant and VSV-sensitive disease, respectively, to investigate the mechanisms underlying resistance to VSV-mediated oncolysis. PC3 cells exhibited elevated baseline expression of multiple pro-survival and anti-apoptotic genes including members of the BCL-2 family and components of the NF-κB signaling pathway, as compared to LNCaP cells. Expression of several pro-apoptotic genes, including BIM, PUMA, and NOXA, was further upregulated upon VSV infection of PC3 cells. These findings suggest that PC3 cells are intrinsically “primed” to resist VSV-mediated killing.

VSV infection of PC3 cells also induced anti-apoptotic signaling, creating competing pro-survival and pro-death programs within the infected cells. We therefore hypothesize that PC3 cells resist VSV-mediated killing because the balance of signaling remains shifted toward survival rather than apoptosis following infection, despite the induction of pro-apoptotic genes. This interpretation is consistent with previous observations showing that PC3 cells remain resistant to VSV-mediated killing despite exposure to viral doses substantially higher than those required to induce cell death in LNCaP cells.

Complementary phospho-proteomic analyses demonstrated activation of both canonical and non-canonical NF-κB signaling, as well as increased PI3K-Akt and MSK1 pathway activity in infected PC3 cells. Together, these findings support a model in which NF-κB-centered signaling networks cooperate with PI3K-Akt and MSK1 pathways to promote cell survival and maintain resistance to VSV-induced apoptosis.

The PI3K-Akt pathway is among the most frequently dysregulated signaling pathways in cancer and plays a well-established role in tumor progression, proliferation, and survival [22]. Consequently, considerable effort has been directed toward targeting this pathway therapeutically [23]. Our findings suggest that targeting downstream anti-apoptotic effectors may represent an alternative or complementary strategy for overcoming resistance to oncolytic virotherapy. Pharmacological inhibition of BCL-2 family proteins restored sensitivity of PC3 cells to VSV-mediated killing, supporting the importance of anti-apoptotic signaling in the resistant phenotype. Similarly, inhibition of NF-κB signaling sensitized PC3 cells to viral oncolysis, further implicating this pathway as a key contributor to resistance.

While these findings highlight the therapeutic potential of combining VSV with inhibitors of pro-survival signaling pathways, several considerations remain. BCL-2 family inhibitors are potent inducers of apoptosis and may be associated with dose-limiting toxicities, particularly when multiple family members are targeted simultaneously. Therefore, careful optimization of treatment regimens will be necessary to maximize tumor-specific killing while minimizing toxicity to normal tissues. In addition, the precise mechanisms by which NF-κB, PI3K-Akt, and MSK1 signaling cooperate to suppress apoptosis in VSV-infected PC3 cells remain incompletely understood. Future studies should investigate downstream effectors of these pathways, including potential contributions from mTOR signaling and other survival networks.

In conclusion, our findings indicate that resistance to VSV-mediated oncolysis in aggressive prostate cancer cells is associated with constitutive activation of pro-survival signaling pathways and sustained activation of NF-κB-centered signaling networks following infection. Pharmacological inhibition of these pathways restores sensitivity to viral killing, supporting their functional role in the resistant phenotype. These results provide mechanistic insight into resistance to oncolytic virotherapy and support the development of combination therapeutic approaches designed to enhance the efficacy of VSV-based treatments for prostate cancer.

