## Supplementary for "Multi-omics Profiling Reveals an NF-κB-Driven Anti-apoptotic Network Underlying Resistance to Oncolytic VSV in Prostate Cancer Cells"

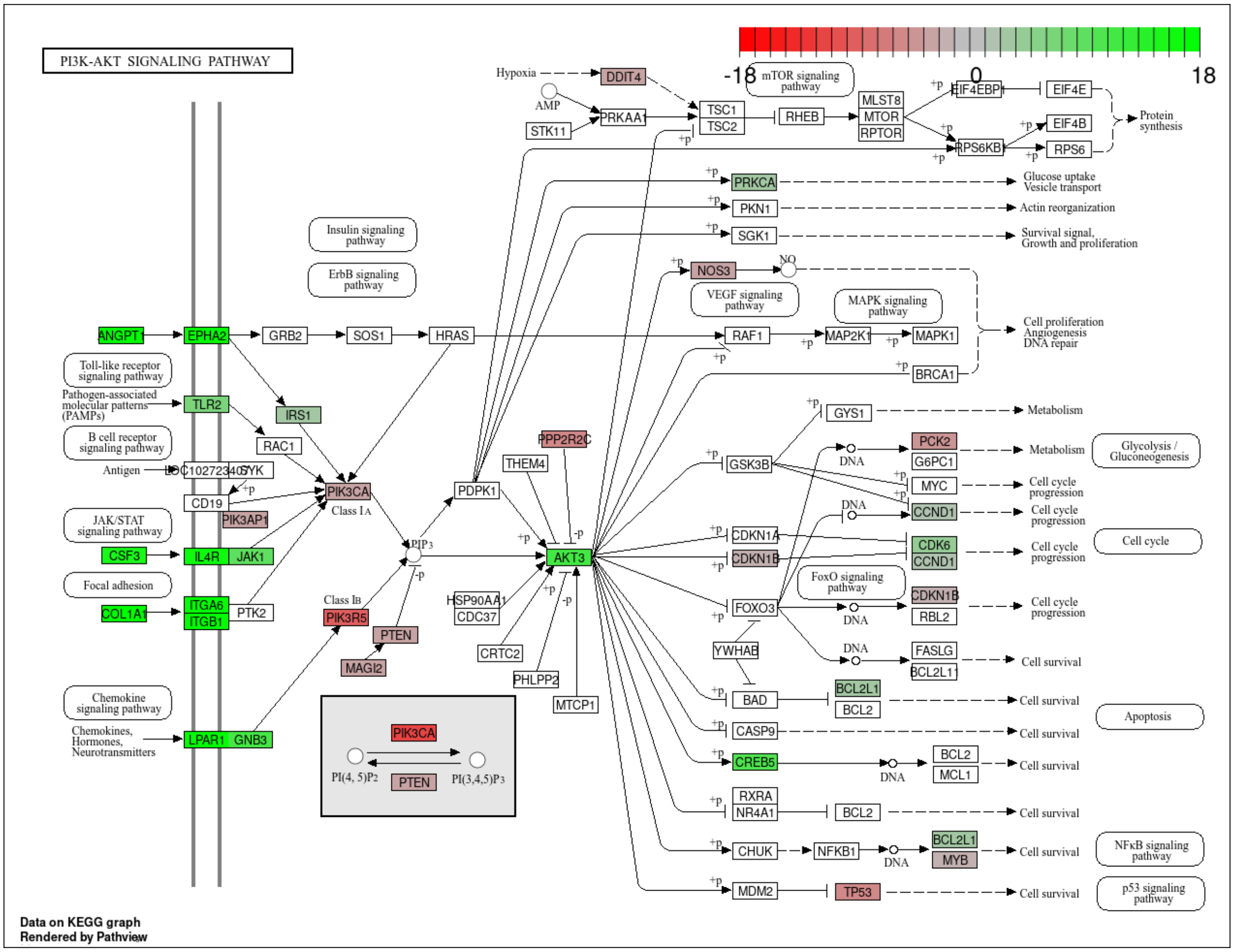


Figure S.1. PC3 cells express higher levels of many pro-survival genes in the absence of viral infection, as compared to LNCaP cells. KEGG pathway enrichment analysis of the PI3K-AKT pathway in uninfected PC3 cells as compared to LNCaP. The color scale reflects log2 Fold Changes (green = upregulated in PC3 cells, gray = unchanged, red = downregulated). The Bioconductor package pathview (v.1.52.0) was used to generate this figure in R.

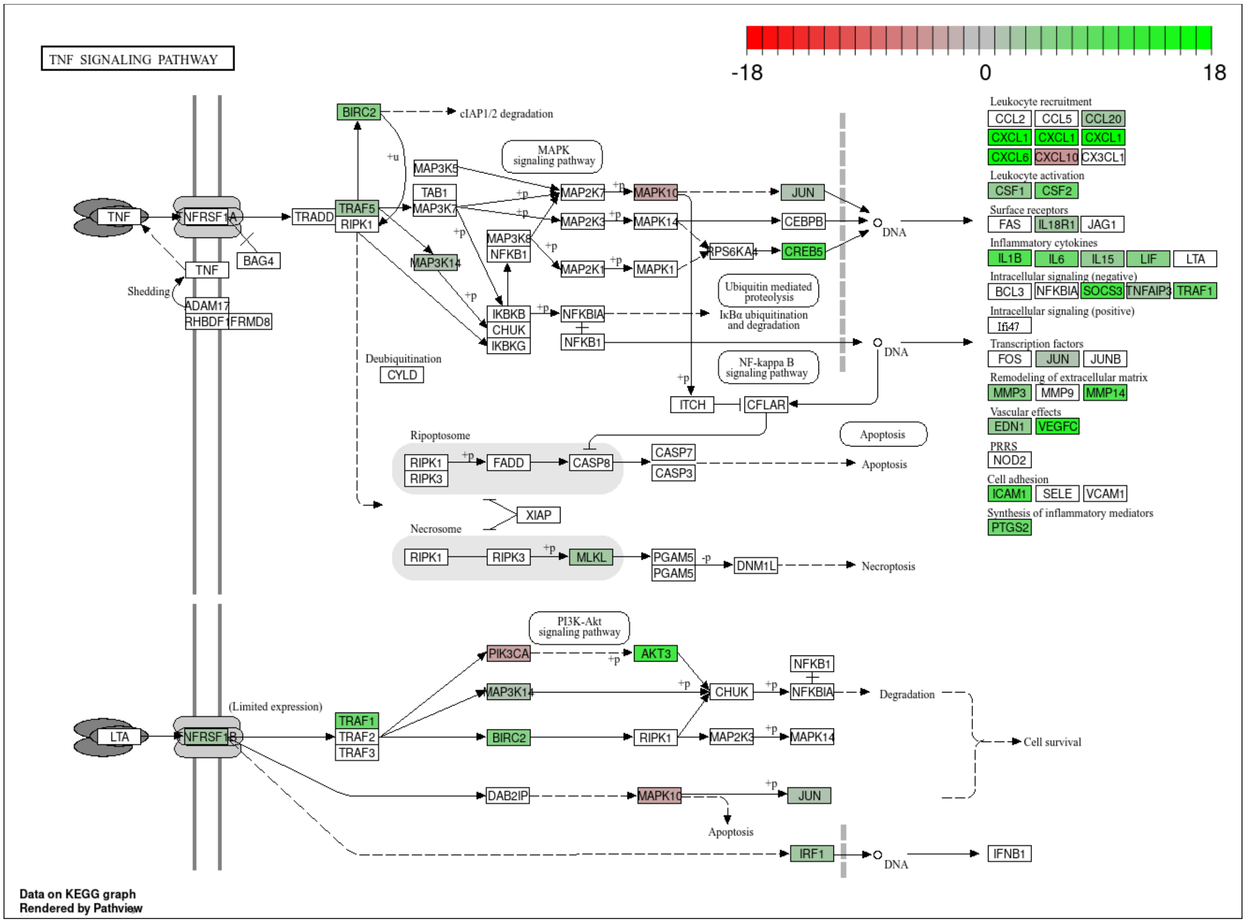


Figure S.2. PC3 cells express higher levels of many pro-survival genes in the absence of viral infection, as compared to LNCaP cells. KEGG pathway enrichment analysis of the TNF pathway in uninfected PC3 cells as compared to LNCaP. The color scale reflects log2 Fold Changes (green = upregulated in PC3 cells, gray = unchanged, red = downregulated). The Bioconductor package pathview (v.1.52.0) was used to generate this figure in R.

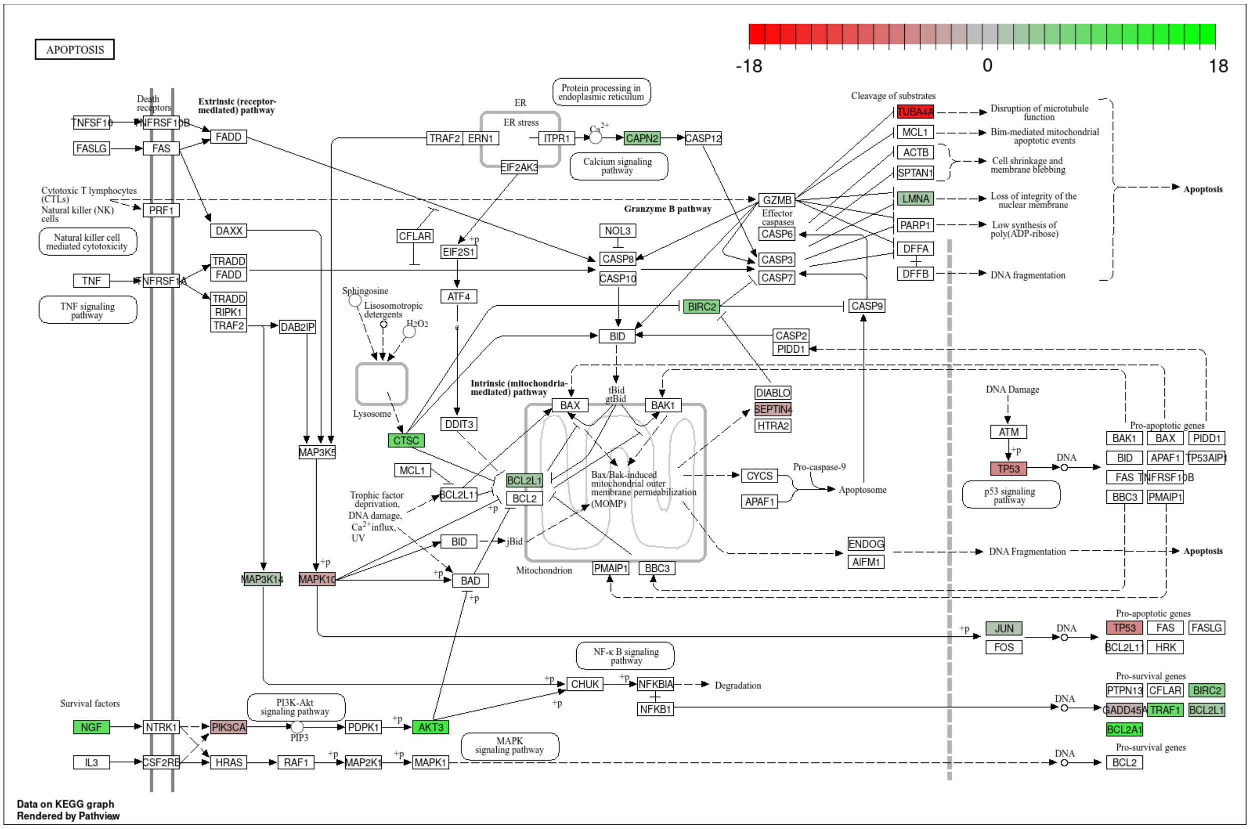


Figure S.3PC3 cells express higher levels of many pro-survival genes in the absence of viral infection, as compared to LNCaP cells. KEGG pathway enrichment analysis of the Apoptosis pathway in uninfected PC3 cells as compared to LNCaP. The color scale reflects log2 Fold Changes (green = upregulated in PC3 cells, gray = unchanged, red = downregulated).. The Bioconductor package pathview (v.1.52.0) was used to generate this figure in R.

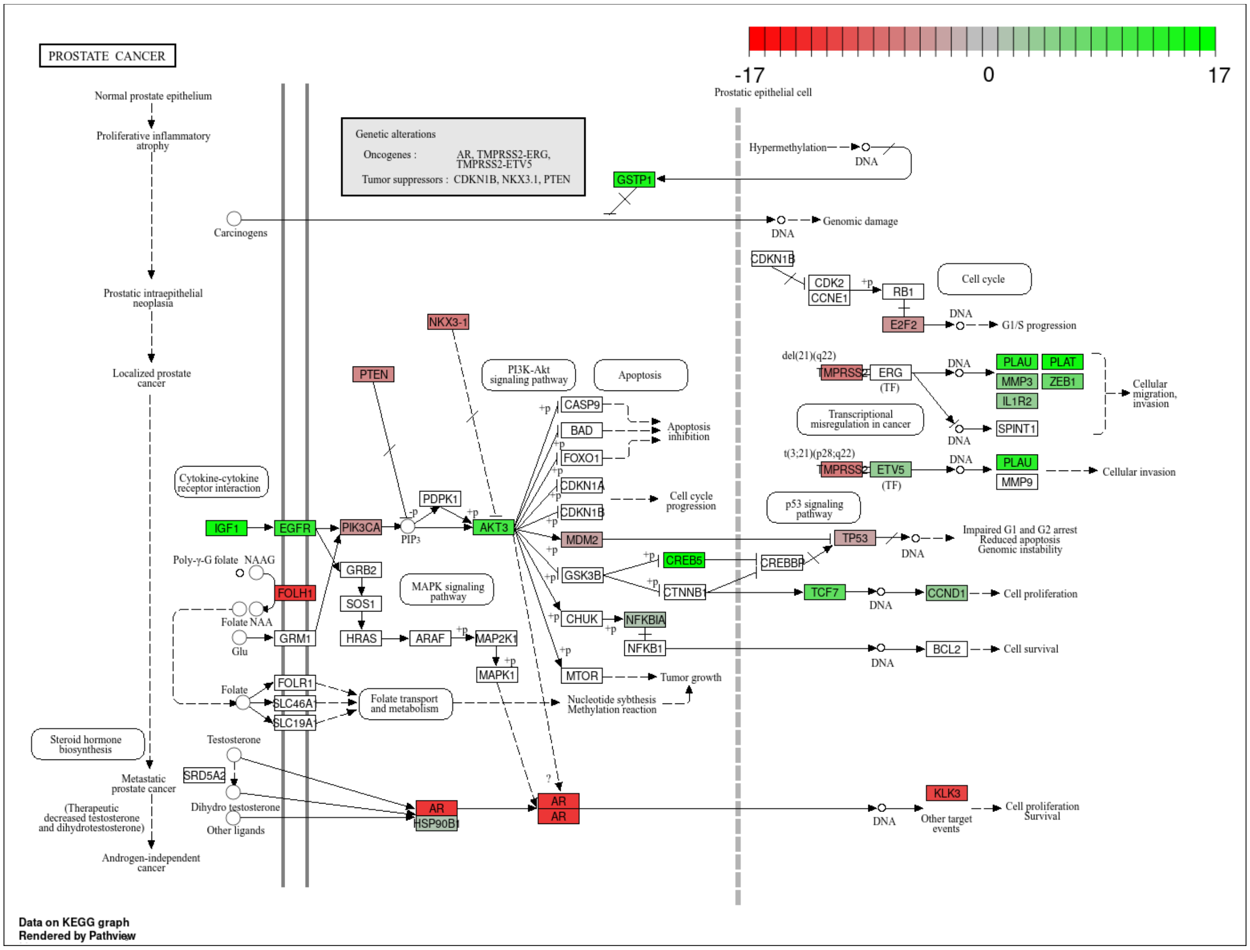


Figure S.4. PC3 cells express higher levels of many pro-survival genes in the absence of viral infection, as compared to LNCaP cells. KEGG pathway enrichment analysis of the Prostate Cancer Signaling pathway in uninfected PC3 cells as compared to LNCaP. The color scale reflects log2 Fold Changes (green = upregulated in PC3 cells, gray = unchanged, red = downregulated). The Bioconductor package pathview (v.1.52.0) was used to generate this figure in R.

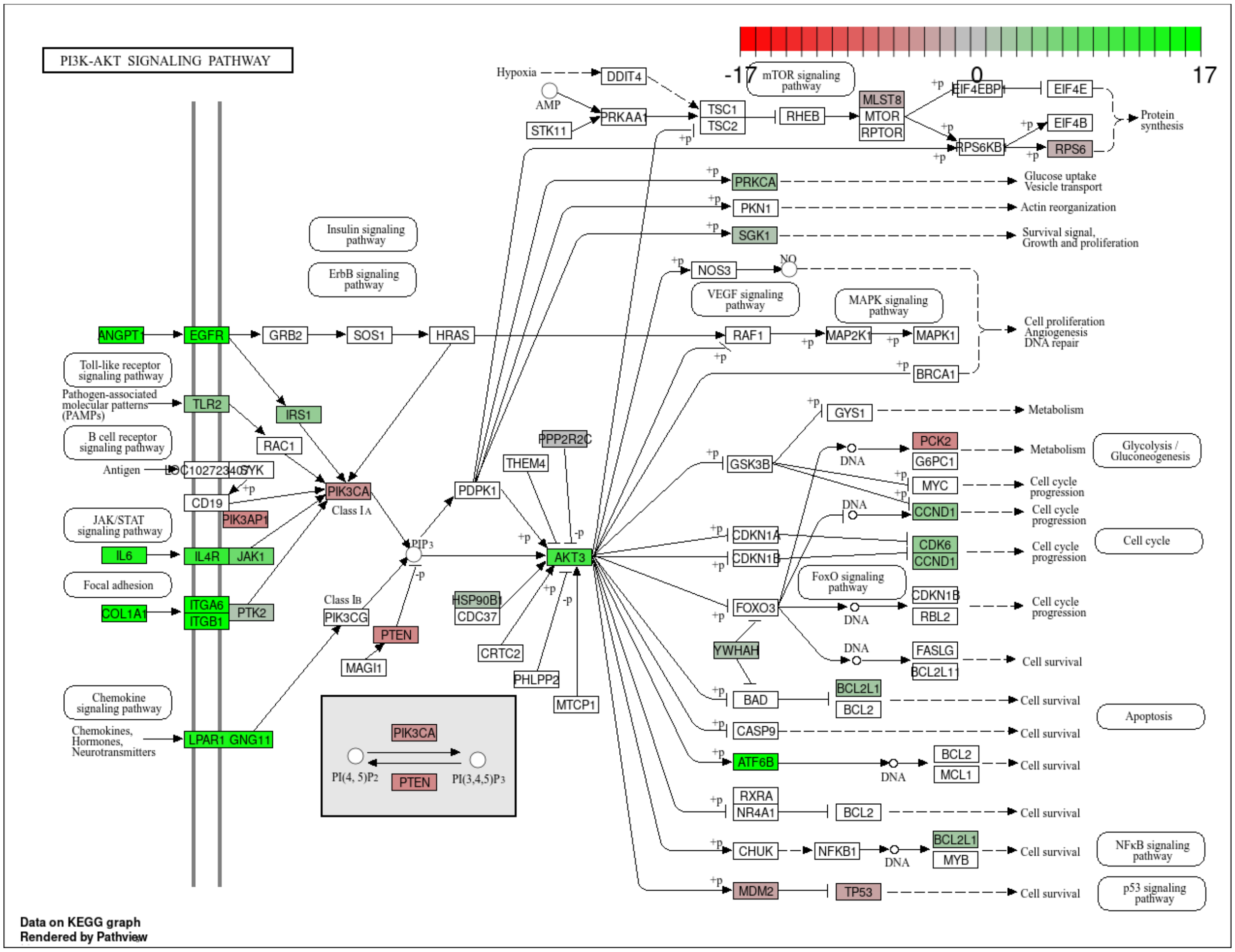


Figure S.5. VSV infection causes upregulation of many pro-survival and antiviral genes in PC3 cells, as compared to LNCaP cells. KEGG PI3K-AKT Signaling pathway enrichment analysis in VSV-infected PC3 cells compared to infected LNCaP cells. The color scale reflects log2 Fold Changes (green = upregulated in PC3 cells, gray = unchanged, red = downregulated). The Bioconductor package pathview (v.1.52.0) was used to generate this figure in R.

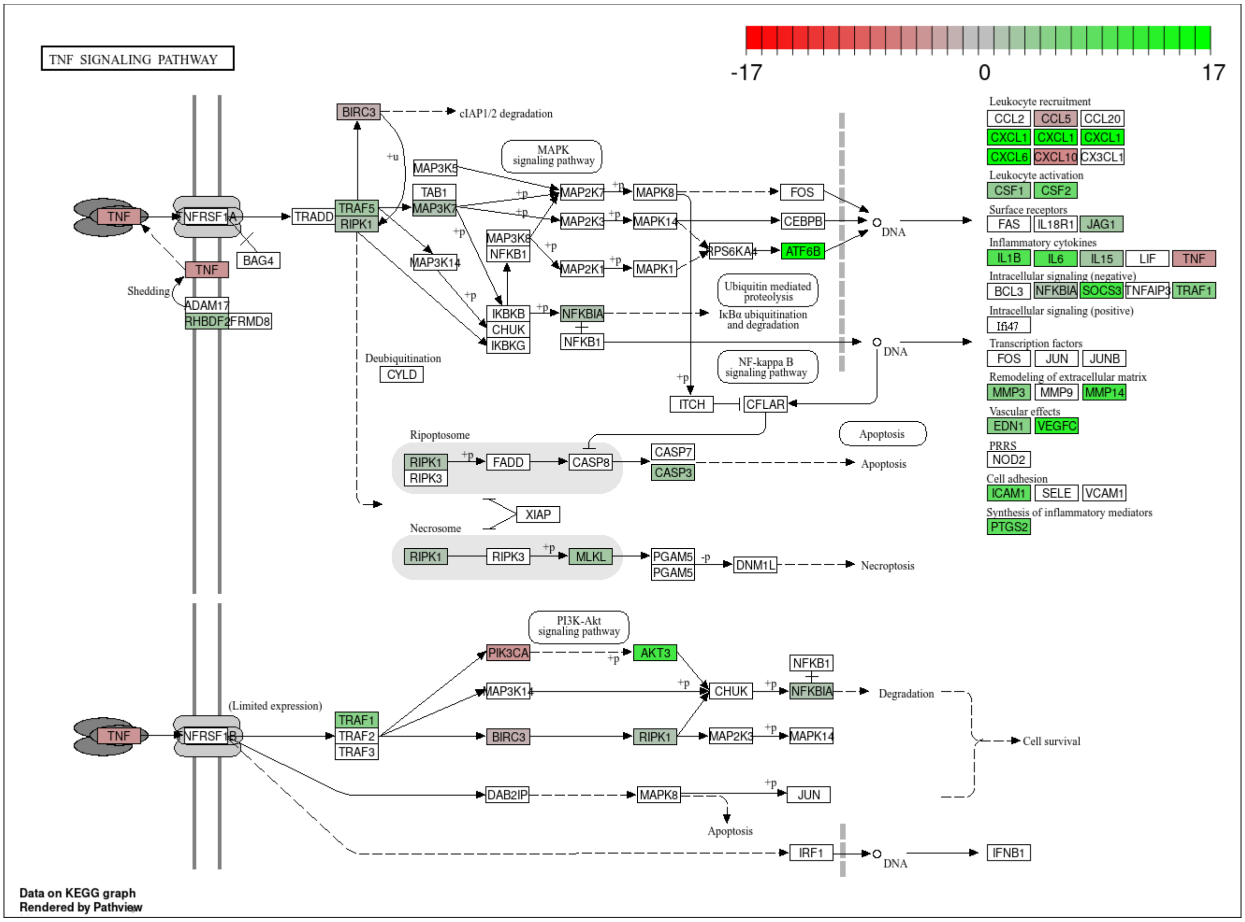


Figure S.6. VSV infection causes upregulation of many pro-survival and antiviral genes in PC3 cells, as compared to LNCaP cells. KEGG TNF Signaling pathway enrichment analysis in VSV-infected PC3 cells compared to infected LNCaP cells. The color scale reflects log2 Fold Changes (green = upregulated in PC3 cells, gray = unchanged, red = downregulated). The Bioconductor package pathview (v.1.52.0) was used to generate this figure in R.

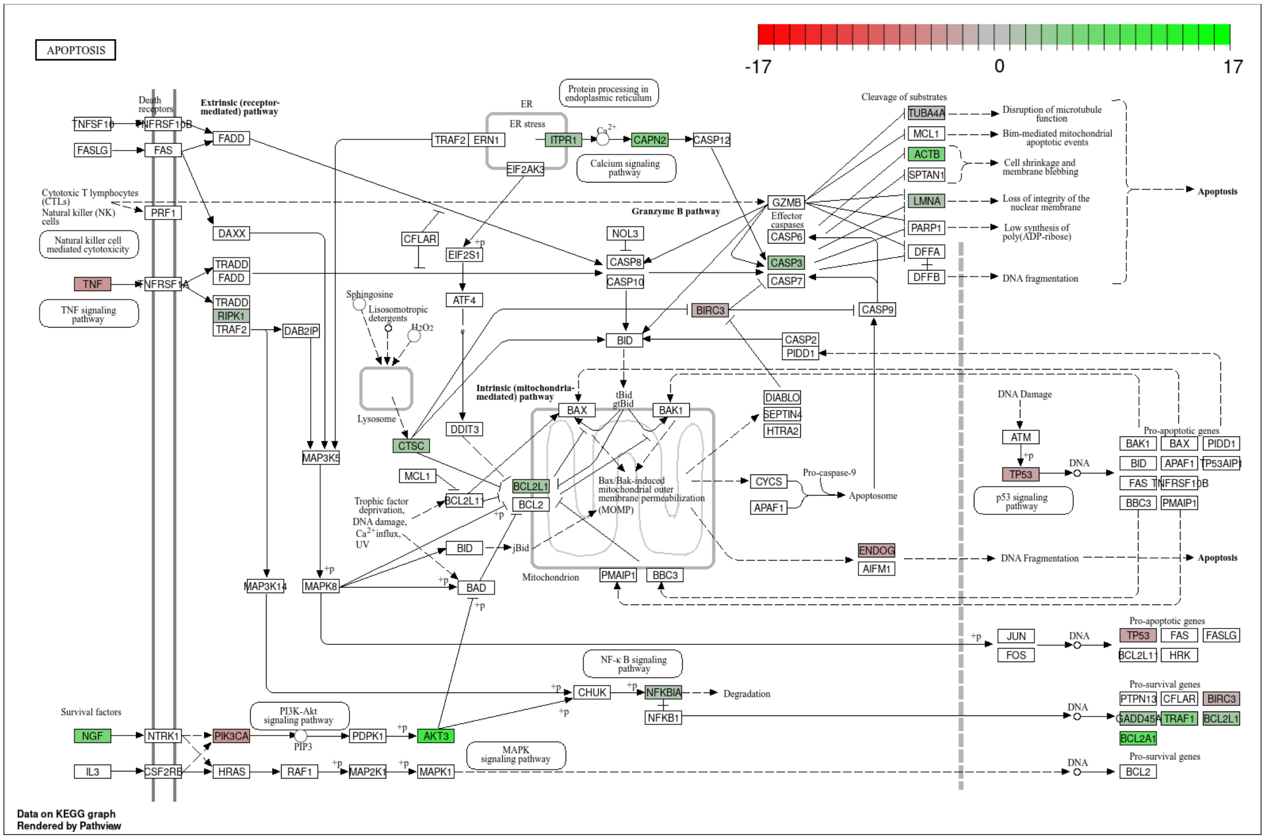


Figure S.7. VSV infection causes upregulation of many pro-survival and antiviral genes in PC3 cells, as compared to LNCaP cells. KEGG Apoptosis Signaling pathway enrichment analysis in VSV-infected PC3 cells compared to infected LNCaP cells. The color scale reflects log2 Fold Changes (green = upregulated in PC3 cells, gray = unchanged, red = downregulated). The Bioconductor package pathview (v.1.52.0) was used to generate this figure in R.

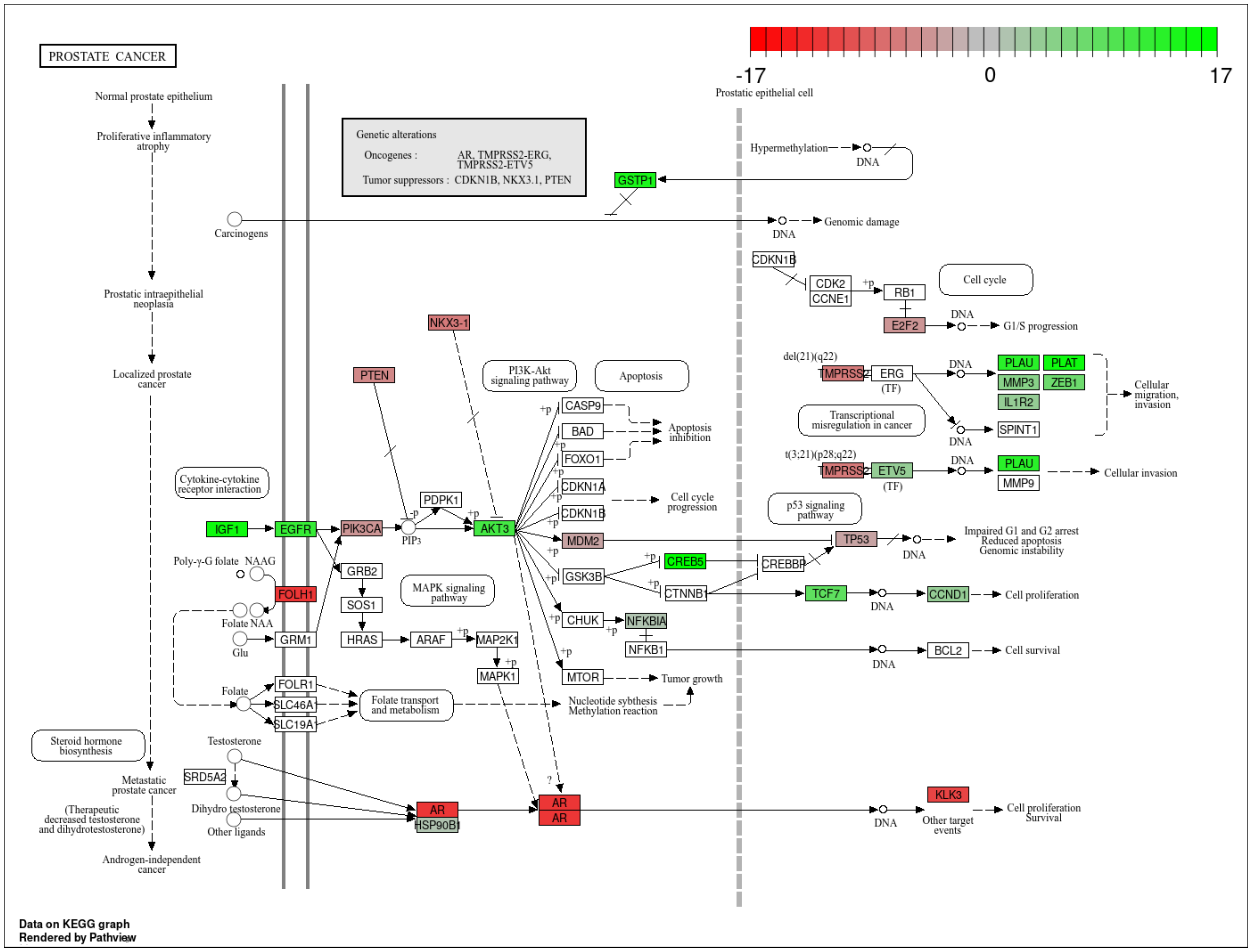


Figure S.8. VSV infection causes upregulation of many pro-survival and antiviral genes in PC3 cells, as compared to LNCaP cells. KEGG prostate cancer Signaling pathway enrichment analysis in VSV-infected PC3 cells compared to infected LNCaP cells. The color scale reflects log2 Fold Changes (green = upregulated in PC3 cells, gray = unchanged, red = downregulated). The Bioconductor package pathview (v.1.52.0) was used to generate this figure in R.

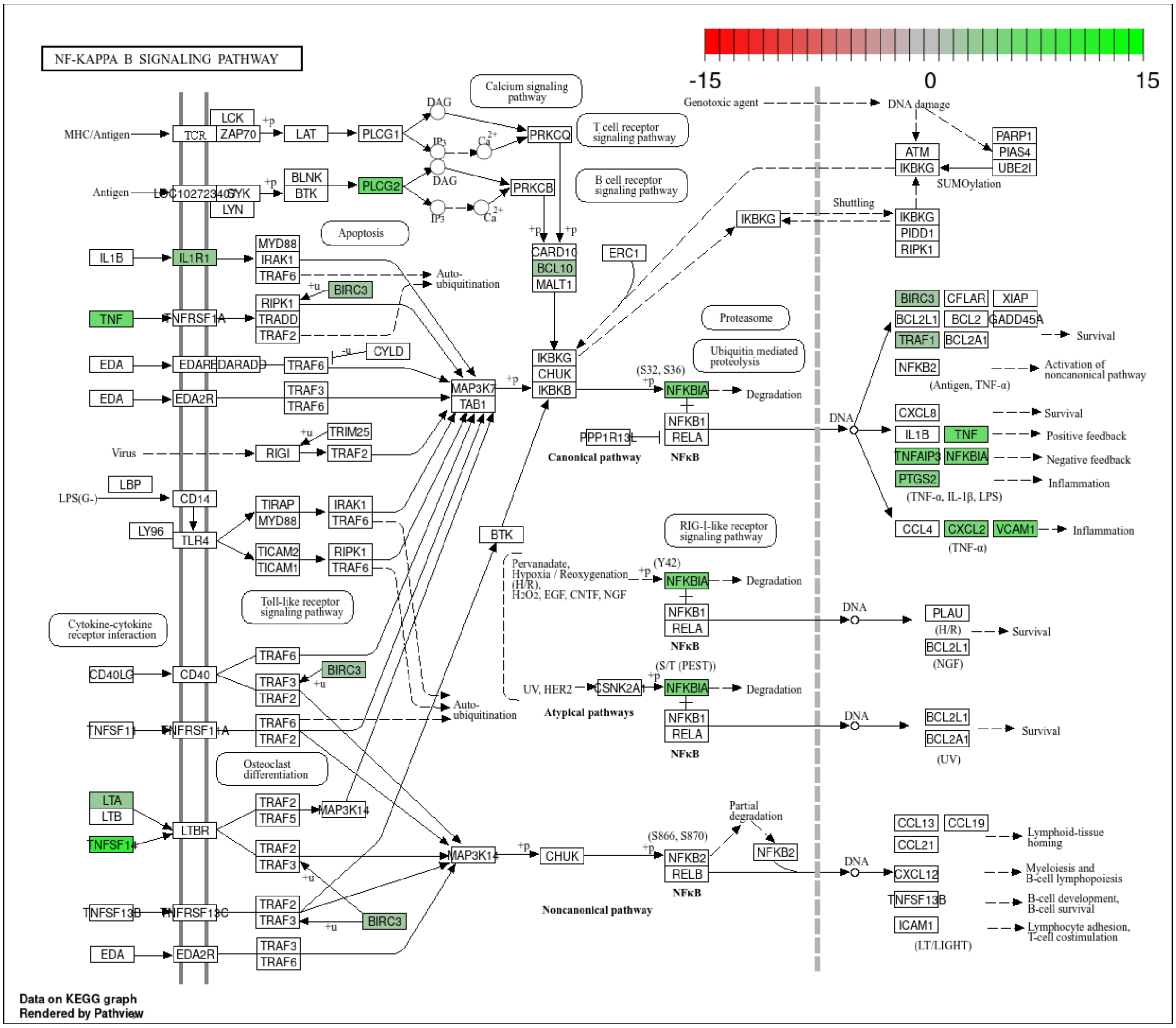


Figure S.9. VSV infection causes upregulation of many pro-survival and antiviral genes in infected PC3 cells, as compared to their Mock. KEGG NF-κB Signaling pathway enrichment analysis in VSV-infected PC3 cells compared to uninfected PC3. The color scale reflects log2 Fold Changes (green = upregulated in PC3 cells, gray = unchanged, red = downregulated). The Bioconductor package pathview (v.1.52.0) was used to generate this figure in R.

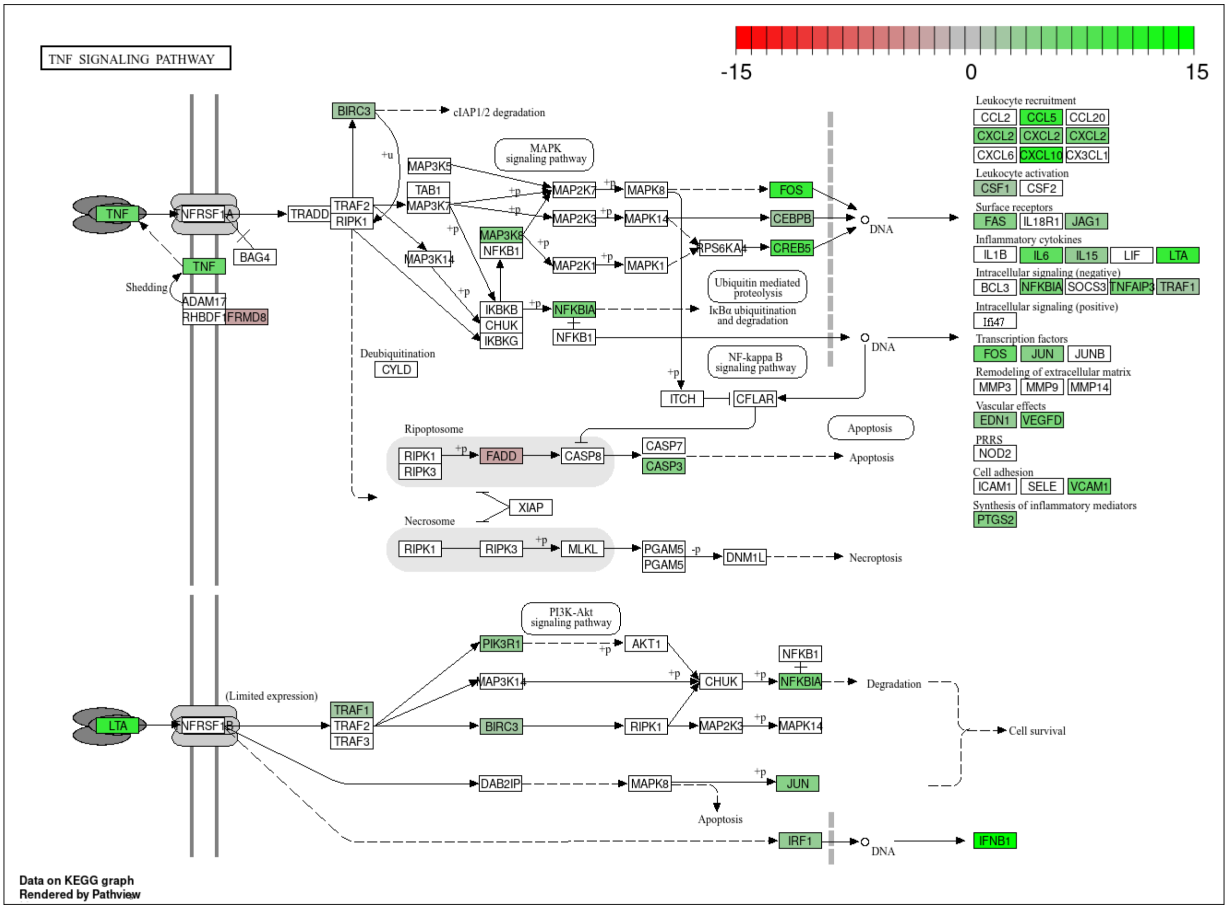


Figure S.10. VSV infection causes upregulation of many pro-survival and antiviral genes in infected PC3 cells, as compared to their Mock. KEGG TNF Signaling pathway enrichment analysis in VSV-infected PC3 cells compared to uninfected PC3. The color scale reflects log2 Fold Changes (green = upregulated in PC3 cells, gray = unchanged, red = downregulated). The Bioconductor package pathview (v.1.52.0) was used to generate this figure in R.
